# Improving the annotation of the *Heterorhabditis bacteriophora* genome

**DOI:** 10.1101/225904

**Authors:** Florence McLean, Duncan Berger, Dominik R. Laetsch, Hillel T. Schwartz, Mark Blaxter

## Abstract

Genome assembly and annotation remains an exacting task. As the tools available for these tasks improve, it is useful to return to data produced with earlier instances to assess their credibility and correctness. The entomopathogenic nematode *Heterorhabditis bacteriophora* is widely used to control insect pests in horticulture. The genome sequence for this species was reported to encode an unusually high proportion of unique proteins and a paucity of secreted proteins compared to other related nematodes. We revisited the *H. bacteriophora* genome assembly and gene predictions to ask whether these unusual characteristics were biological or methodological in origin. We mapped an independent resequencing dataset to the genome and used the blobtools pipeline to identify potential contaminants. While present (0.2% of the genome span, 0.4% of predicted proteins), assembly contamination was not significant. Re-prediction of the gene set using BRAKER1 and published transcriptome data generated a predicted proteome that was very different from the published one. The new gene set had a much reduced complement of unique proteins, better completeness values that were in line with other related species’ genomes, and an increased number of proteins predicted to be secreted. It is thus likely that methodological issues drove the apparent uniqueness of the initial *H. bacteriophora* genome annotation and that similar contamination and misannotation issues affect other published genome assemblies.

## Introduction

The sequencing and annotation of a species’ genome is often but the first step in exploiting these data for comprehensive biological understanding. As with all scientific endeavour, genome sequencing technologies and the bioinformatics toolkits available for assembly and annotation are being continually improved. It should come as no surprise therefore that first estimates of genome sequences and descriptions of the genes they contain can be improved. For example, the genome of the nematode *Caenorhabditis elegans* was the first animal genome to be sequenced [1]. The genome sequence and annotations have been updated many times since, as further exploration of this model organism revealed errors in original predictions, such that today, with release WS260 (http://www.wormbase.org/) [2], very few of the 19099 protein coding genes announced in the original publication [1] retain their original structure and sequence. The richness of the annotation of *C. elegans* is driven by the size of the research community that uses this model species. However for most species, where the community using the genome data is small or less-well funded, initial genome sequences and gene predictions are not usually updated.

*Heterorhabditis bacteriophora* is an entomopathogenic nematode which maintains a mutualistic association with the bacterium *Photorhabdus luminescens.* Unlike many other parasitic nematodes, it is amenable to *in vitro* culture [3] and is therefore of interest not only to evolutionary and molecular biologists investigating parasitic and symbiotic systems, but also to those concerned with the biological control of insect pests [4, 5]. *P. luminescens* colonises the anterior intestine of the free-living infective juvenile stage (IJ). IJs are attracted to insect prey by chemical signals [6, 7]. On contacting a host, the IJs invade the insect’s haemocoel and actively regurgitate *P. luminescens* into the haemolymph. The bacterial infection rapidly kills the insect, and *H. bacteriophora* grow and reproduce within the cadaver. After 2-3 cycles of replication, the nematode progeny develop into IJs, sequester *P. luminescens* and seek out new insect hosts.

Axenic *H. bacteriophora* IJs are unable to develop past the L1 stage [8], and *H. bacteriophora* may depend on *P. luminescens* for secondary metabolite provision [9, 10]. Mutation of the global post-transcriptional regulator Hfq in *P. luminescens* reduced the bacterium’s secondary metabolite production and led to failed nematode development, despite the bacterium maintaining virulence against host (*Galleria mellonella*) larvae [11]. Together these symbionts are efficient killers of pest (and other) insects, and understanding of the molecular mechanisms of host killing could lead to new insecticides.

*H. bacteriophora* was selected by the National Human Genome Research Initiative as a sequencing target [12]. Genomic DNA from axenic cultures of the inbred strain *H. bacteriophora* TTOI was sequenced using Roche 454 technology and a high quality 77 Mb draft genome assembly produced [13]. This assembly was predicted (using JIGSAW [14]) to encode 21250 proteins. Almost half of these putative proteins had no significant similarity to entries in the GenBank non-redundant protein database, suggesting an explosion of novelty in this nematode. The predicted *H. bacteriophora* proteome had fewer orthologues of Kyoto Encyclopedia of Genes and Genomes loci in the majority of metabolic categories than nine other nematodes. *H. bacteriophora* was also predicted to have a relative paucity of secreted proteins compared to free-living nematodes, postulated to reflect a reliance on *P. luminescens* for secreted effectors [13]. The 5.7 Mb genome of *P. luminescens* has also been sequenced [15]. The *H. bacteriophora* proteome had fewer shared orthologues when clustered and compared to other rhabditine (Clade V) nematodes (including *Caenorhabditis elegans* and the many animal parasites of the Strongylomorpha) [16].

In preliminary analyses we noted that while the genome sequence itself had high completeness scores when assessed with the Core Eukaryote Gene Mapping Approach (CEGMA) [17] (99.6% complete) and Benchmarking Universal Single-Copy Orthologs (BUSCO) [18] (80.9% complete and 5.6% fragmented hits for the BUSCO Eukaryota gene set), the predicted proteome scored poorly (47.8% complete and 34.7% fragmented by BUSCO; see below). Another unusual feature of the *H. bacteriophora* gene set was the proportion of non-canonical splice sites (i.e. those with a 5’ GC splice donor site, as opposed to the normal 5’ GT). Most nematode (and other metazoan) genomes have low proportions of non-canonical introns (less than 1%), but the published gene models had over 9% non-canonical introns. This is more than double the proportion predicted for *Globodera rostochiensis*, a plant parasitic nematode where the unusually high proportion of non-canonical introns was validated *via* manual curation [19].

If these unusual characteristics reflect a truly divergent proteome, the novel proteins in *H. bacteriophora* may be crucial in its particular symbiotic and parasitic relationships, and of great interest to development of improved strains for horticulture. However, it is also possible that contamination of the published assembly or annotation artefacts underpin these unusual features. We re-examined the *H. bacteriophora* genome and gene predictions, and used more recent tools to re-predict protein coding genes from the validated assembly. As the BRAKER1 predictions were demonstrably better than the original ones, we explored whether some of the unusual characteristics of the published protein set, in particular the level of novelty and the proportion of secreted proteins, were supported by the BRAKER1 protein set.

## Results

### No evidence for substantial contamination of the *H. bacteriophora* genome assembly

We used BlobTools [20] to assess the published genome sequence [13] for potential contamination. The raw read data from the published assembly was not available on the trace archive or short read archive (SRA). We thus utilised new Illumina short-read re-sequencing data generated from strain G2a1223, an inbred derivative of *H. bacteriophora* strain “Gebre”, isolated by Adler Dillman in Moldova. G2a1223 has about 1 single-nucleotide change per ∼2000 nucleotides compared to the originally-sequenced TT01 strain. G2a1223 was grown in culture on the non-colonising bacterium *Photorhabdus temperata*. The majority of these data (96.3% of the reads) mapped as pairs to the assembly, suggesting completeness of the published assembly with respect to the new raw read data. In addition, 99.96% of the published assembly had at least 10-fold coverage from the new raw reads.

The assembly was explored using a taxon-annotated GC-coverage plot, with coverage taken from the new Illumina data and sequence similarity from the NCBI nucleotide database (nt) (Figure 1). *H. bacteriophora* was excluded from the database search used to annotate the scaffolds to exclude self hits from the published assembly. All large scaffolds clustered congruently with respect to read coverage and CG content. A few (57) scaffolds had best BLASTn matches to phyla other than Nematoda (Table 1). A small amount (5 kb) of likely remaining *P. luminescens* contamination was noted. We identified 100 kb of the genome of a strain of the common culture contaminant bacterium *Stenotrophomonas maltophilia* [21]. Contamination of the assembly with *S. maltophilia* was acknowledged [13] but removal of scaffolds before annotation was not discussed. Two high-coverage scaffolds that derived from the *H. bacteriophora* mitochondrial genome were annotated as “undefined Eukaryota” because of taxonomic misclassification in the NCBI nt database. Many scaffolds with coverages close to that of the expected nuclear genome had best matches to two unexpected sources: the platyhelminths *Echinostoma caproni* and *Dicrocoelium dendriticum,* and several hymenopteran arthropods. Inspection of these matches showed that they were due to high sequence similarity to a family of *H. bacteriophora* mariner-like transposons [22] and thus these were classified as *bona fide* nematode nuclear sequences. A group of scaffolds contained what appears to be a *H. bacteriophora* nuclear repeat with highest similarity to histone H3.3 sequences from Diptera and Hymenoptera. The remaining scaffolds had low-scoring nucleotide matches to a variety of chordate, chytrid and arthropod sequences from deeply conserved genes (tubulin, kinases), but had coverages similar to other nuclear sequences.

**Figure 1.**
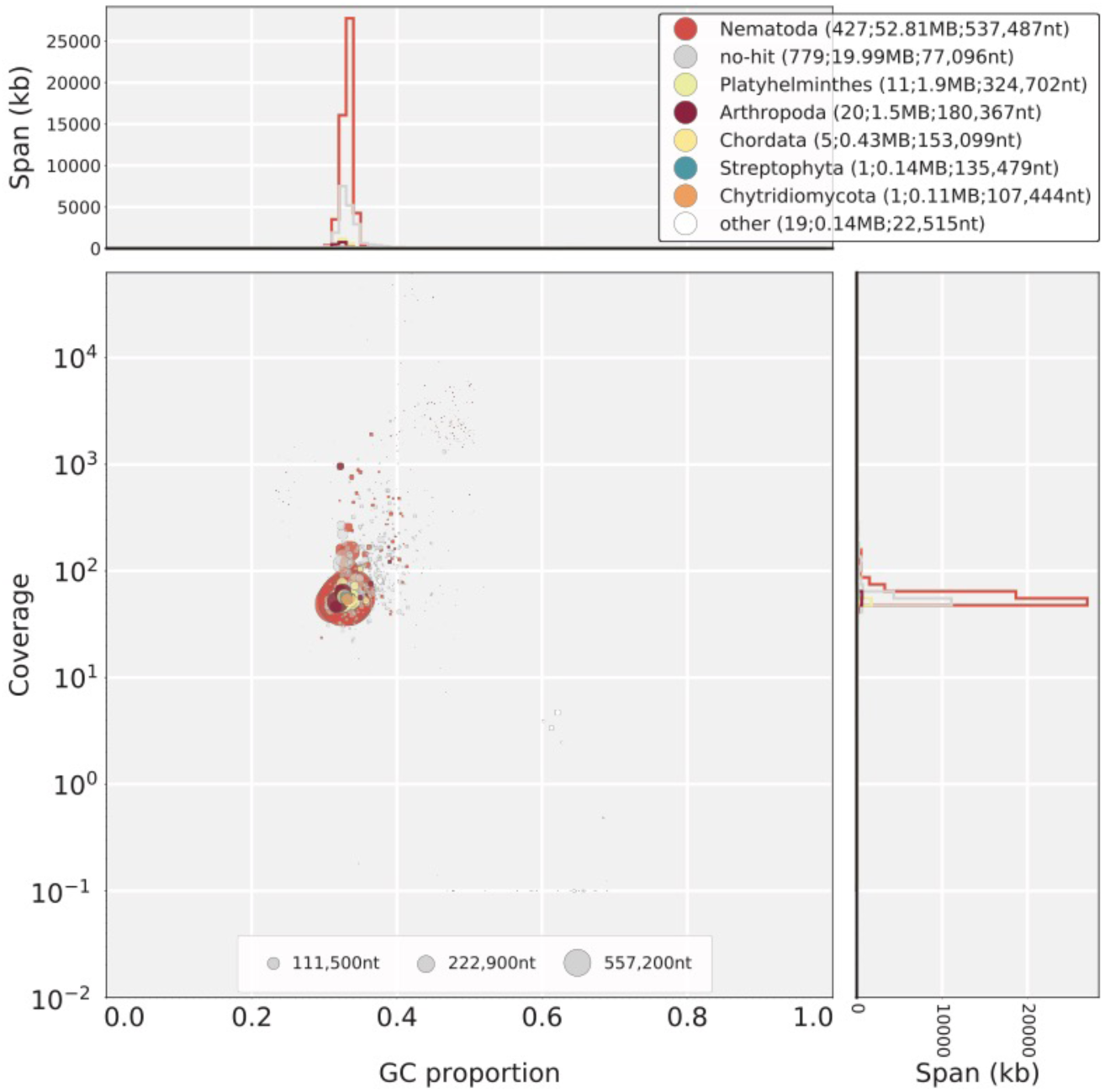
Taxon-annotated GC-coverage plot of the *H. bacteriophora* assembly. Bottom left panel: Each scaffold or contig is represented by a single filled circle. Each scaffold is placed in the main panel based on its GC proportion (X axis) and coverage by reads from the Illumina re-sequencing project (Y axis). The fill colour of the circle indicates the taxon of the top BLASTn hit in the NCBI nt database for that scaffold. The colours are annotated in the top right hand key, which indicates taxon assignment and (in brackets) the number of contigs and scaffolds so assigned, their total span, and their N50 length. The circles are scaled to scaffold length, as indicated in the key at the base of the main panel. Right panel:Nucleotide span in kb at each coverage level. Top panel:Nucleotide span in kb at each GC proportion.

**Table 1.**
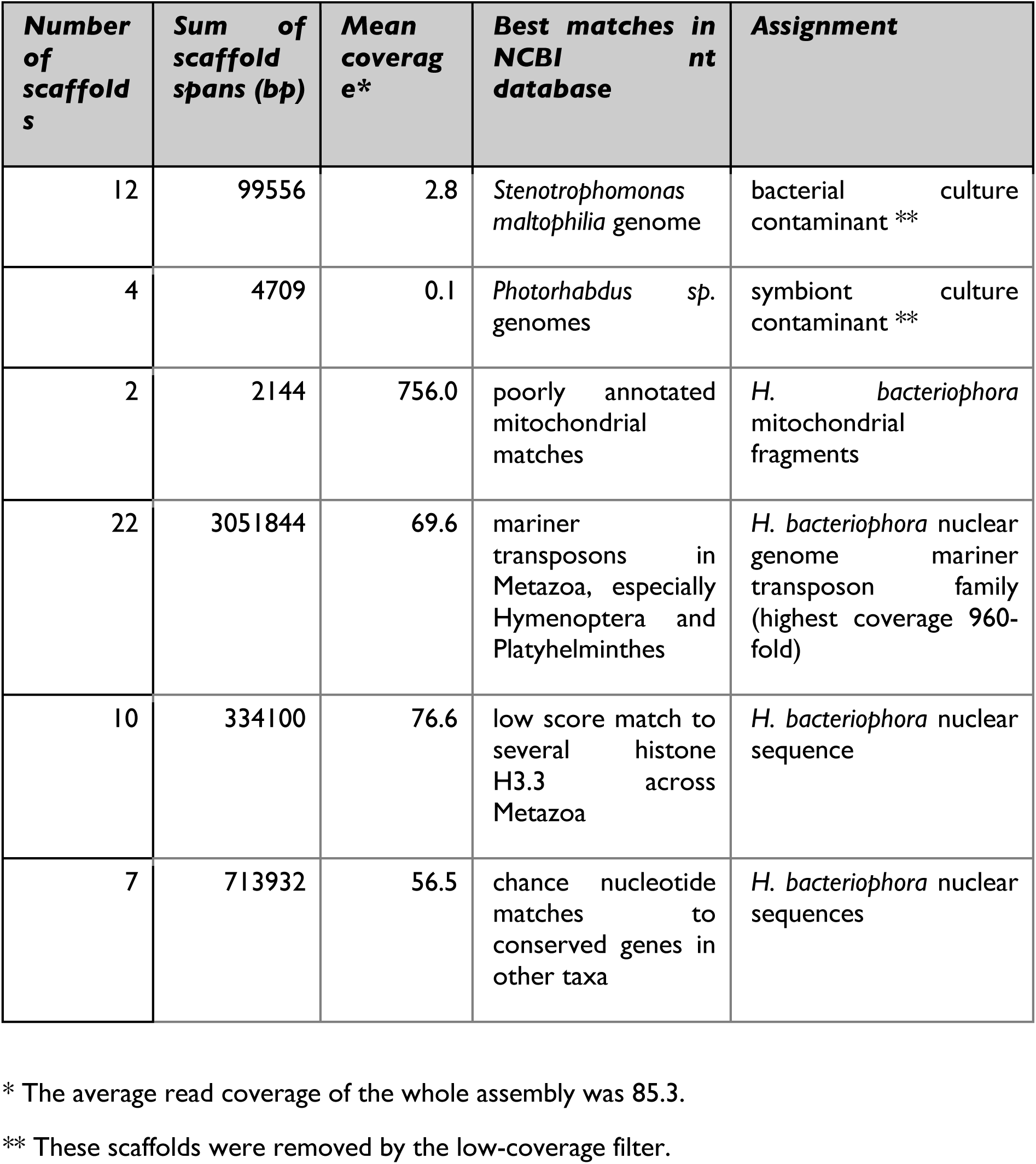
Contamination screening of the *H. bacteriophora* assembly

Scaffolds with average coverage of less than 10-fold were removed from the assembly (35 scaffolds spanning 132949 bases, 0.2% of the total span; see Supplementary File 1). This removed all scaffolds aligning to *S. maltophilia* and to *Photorhabdus* spp. (104 kb). The origins of the additional 28 kb were not investigated. In the published annotation [13], 76 genes were predicted from these scaffolds.

#### Improved gene predictions are biologically credible and have unexceptional novelty

New gene predictions were generated from a soft-masked version of the filtered assembly using the RNA-seq based annotation pipeline BRAKER1 [23], generating 16070 protein predictions from 15747 protein coding genes (see Supplementary File 2). We compared the soft-masked predictions to those from the published analysis [13] (Figure 2, Table 2). The predicted proteins from the new BRAKER1/soft-masked gene set were, on average, longer (Figure 2A). While the average number of introns per gene was the same in the BRAKER1/soft-masked and published predictions, the BRAKER1/soft-masked gene set had more single-exon genes (Figure 2B). Hard masking of the genome and re-prediction resulted in fewer single exon genes, suggesting that many of these putative genes could be derived from repetitive sequence (Supplementary Files 3 and 4), but only 316 of the single exon genes from the BRAKER1/soft-masked assembly had similarity to transposases or transposons. The BRAKER1/soft-masked annotations were taken forward for further analysis.

**Figure 2.**
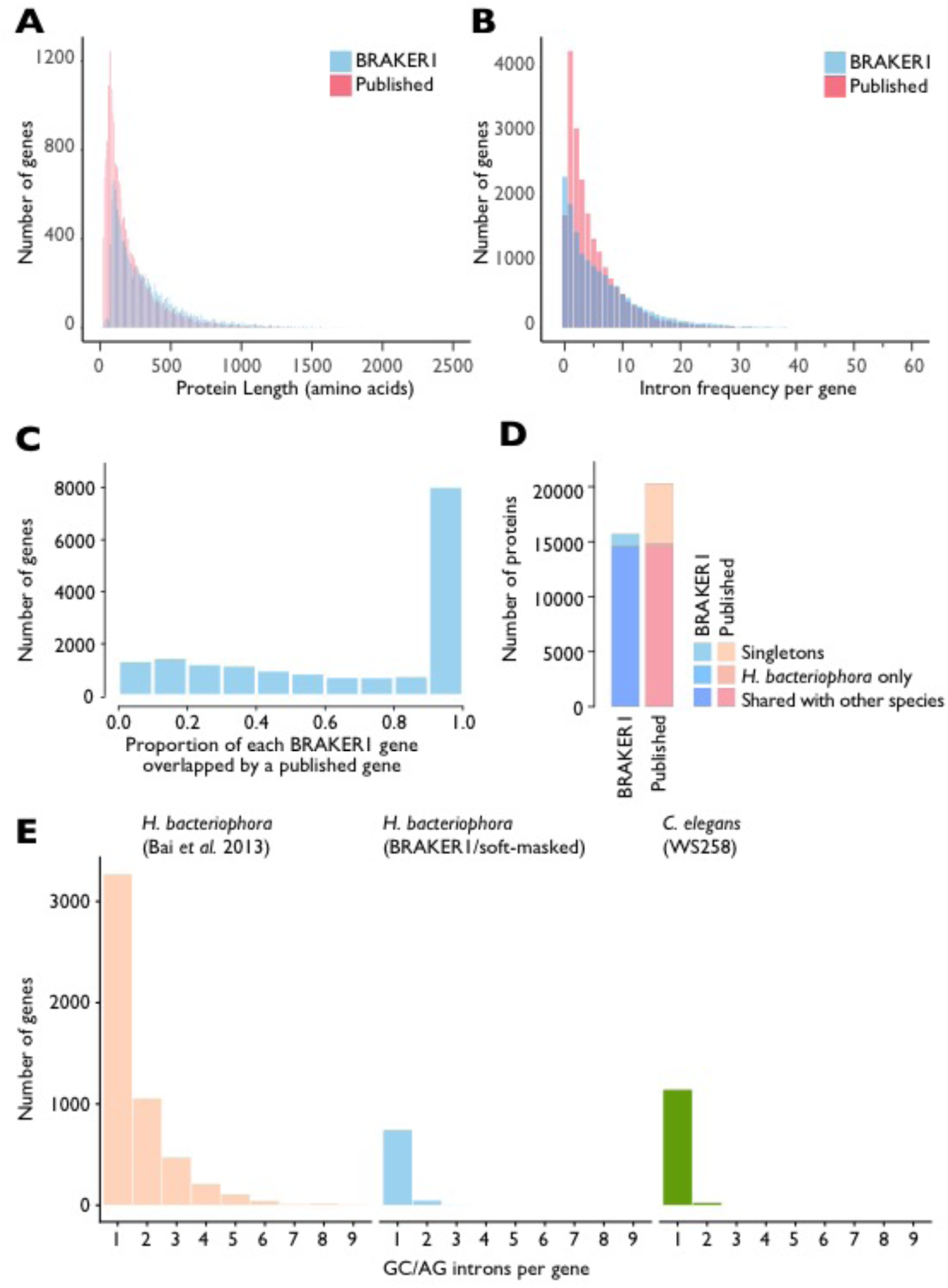
Comparisons of BRAKER1/soft-masked and original gene predictions from *H. bacteriophora*. (A, B) Frequency histograms of intron count (A) and protein length (B) in BRAKER1/soft-masked (blue) and published (yellow) protein coding gene predictions. Outlying proteins longer than >2500 amino-acids (n=40) or genes containing >60 introns (n=20) are not shown. (C) Frequency histogram of the proportion of each BRAKER1 gene prediction overlapped by a published gene prediction at the nucleotide level. (D) Comparison of singleton, proteome-specific, and shared proteins in the published and BRAKER1/soft-masked protein sets. (E) Counts of non-canonical GC/AG introns in gene predictions from the published and BRAKER1 *H. bacteriophora* gene sets, and the model nematode *Caenorhabditis elegans* (WS258). Counts are of genes containing at least one non-canonical GC/AG intron with the specified number of non-canonical introns.

**Table 2.** Comparison of the published and BRAKER1/soft-masked protein coding gene predictions.

| *Prediction set* | *Published [13]* | *BRAKER1/soft-masked* |
| --- | --- | --- |
| Number of protein coding genes predicted | 20964 | 15747 |
| Mean protein length (amino acids) | 218.8 | 344.5 |
| Number of single exon genes | 1728 | 2326 |
| Mean number of exons per gene* | 5.9 | 7.8 |
| Proportion of non-canonical (GC-AG) introns | 8.87% | 0.79% |
| Percentage mapping to publicly available transcriptome reads |  |  |
| *Sanger ESTs* | 80.45% | 84.26% |
| *Roche 454 reads* | 37.18% | 58.03% |
| BUSCO score for proteome |  |  |
| *Complete* | 47.8% | 94% |
| *Fragmented* | 34.7% | 4.3% |
| Number of proteins with no hits in Uniref90 | 8,962 | 2,889 |
| Protein singletons in clustering | 5442 | 1112 |
| Conserved, single-copy orthologues** |  |  |
| *Total* | 2089 | 2330 |
| *Missing* | 377 | 141 |
| *Expanded* | 184 | 84 |

\* Number of exons: number of coding DNA sequence (CDS) entries per gene for BRAKER1 predictions. CDS features, not exons are outputted by AUGUSTUS in general feature format (GFF).
\*\* The list of strict one-to-one orthologues was augmented with protein clusters where 75% of species had single copy representatives (“fuzzy-1-to-1” orthologues identified by KinFin).

Four-fifths (83.3%) of the published protein-coding gene predictions [13] overlapped to some extent with the BRAKER1/soft-masked predictions at the genome level, with a mean of 67% of the nucleotides of each BRAKER1/soft-masked gene covered by a published gene (Figure 2C). Half (8061) of the 15747 BRAKER1/soft-masked gene predictions had an overlap proportion of ≥ 0.9 with the published predictions. At the level of protein sequence only 836 proteins were identical between the two predictions, and only 2099 genes had identical genome start and stop positions.

The BRAKER1/soft-masked and published gene sets were checked for completeness using BUSCO [18], based on the Eukaryota lineage gene set, and *Caenorhabditis* as the species parameter for orthologue finding. The BRAKER1/soft-masked gene set contained a substantially higher percentage of complete, and lower percentage of fragmented BUSCO genes than the published set (Table 2). Two *H. bacteriophora* transcriptome datasets, publicly available Roche 454 data and Sanger expressed sequence tags, were mapped to the published and BRAKER1/soft-masked transcriptomes to assess gene set completeness. This suggested that the BRAKER1/soft-masked transcriptome predictions were more complete than the original (Table 2).

Nearly half (9893/20964; 47.2%) of the published proteins were reported to have no significant matches in the NCBI non-redundant protein database (nr) [13]. This surprising result could be due to a paucity of data from species closely related to *H. bacteriophora* in the NCBI nr database at the time of the search, or inclusion of poor protein predictions in the published set, or both. Targeted investigation of these 9893 orphan proteins here was not possible due to inconsistencies in gene naming in the publically available files. The published and BRAKER1/soft-masked proteomes were compared to the Uniref90 database [24], using DIAMOND v0.9.5 [25] with an expectation value cut-off of Ie^-5^. In the published proteome, 8962 proteins (42.7%) had no significant matches in Uniref90. Thus a relatively poorly populated database was not the main driver for the high number of orphan proteins reported in the published proteome. In the BRAKER1/soft-masked proteome, only 2889 proteins (18.3%) had no hits in the Uniref90 database (Table 2).

OrthoFinder vI.I.4 [26] was used to define orthologous groups in the proteomes of 23 rhabditine (Clade V) nematodes (Supplementary Files 5 and 6) and just the published *H. bacteriophora* protein-coding gene predictions, or just the BRAKER/soft-masked proteome, or both. All proteins <30 amino-acids long were excluded from clustering (see Supplementary File 5). We identified 5442 singletons (26.8% of the proteome) when the analysis included only the published *H. bacteriophora* protein set. An additional 248 proteins formed *H. bacteriophora-specific* orthogroups. Orthology analysis including only the BRAKER/soft-masked protein set predicted 1112 *H. bacteriophora* singletons (7.1% of the proteome) with 167 proteins in *H. bacteriophora-specific* orthogroups (Figure 2D). In comparison, when the orthology analysis included the BRAKER1/soft-masked predictions there were 1858 *C. elegans* singletons (9.2% of the *C. elegans* proteome). Very few universal, single copy orthologues were defined in either analysis. Exploring “fuzzy-1-to-1” orthogroups (where true 1-to-1 orthology was found for greater than 75% of the 24 species - i.e. 18 or more species), the published protein predictions had more missing fuzzy-1-to-1 orthologues than did the BRAKER1/soft-masked predictions (Table 2). In the clustering that included both proteomes, 2019 clusters contained more proteins from the BRAKER1/soft-masked than the published proteome, whereas 2714 contained a larger number contributed from the published than the BRAKER1/soft-masked proteome (Supplementary File 6).

The published *H. bacteriophora* gene set had additional peculiarities. The published set of gene models included 102274 introns, 9069 of which (8.9%) had non-canonical splice sites (i.e. 5’ GC – AG 3’). Some of the genes in the published gene set had up to nine noncanonical introns (Figure 2E). In the BRAKER1/soft-masked gene set there were 109767 introns, 868 (0.8%) of which had non-canonical splice sites. This proportion is in keeping with that found in most other rhabditine nematodes. For example, the extensively manually annotated *C. elegans* has 2429 (0.6%) non-canonical (5’ GC – AG 3’) introns. In *C. elegans* non-canonical introns are frequently found only in alternately spliced, and shorter isoforms, and over 93-99% were in genes that had homologues in other species, depending on the species used in the protein orthology clustering. However, in the published *H. bacteriophora* gene set, 34-49% of the genes with GC – AG introns were in *H. bacteriphora-unique* proteins.

A supermatrix maximum likelihood phylogeny was generated from the fuzzy-1-1 orthologues in the clustering that included both *H. bacteriophora* proteomes (Figure 3; see Supplementary File 7). The phylogeny, rooted with *Pristionchus* spp., shows the *H. bacteriophora* proteomes as sisters. However the BRAKER1/soft-masked proteome has a shorter branch length to *Heterorhabditis’* most recent common ancestor with other Clade V nematodes, suggesting that the published proteome includes uniquely divergent sequences.

**Figure 3.**
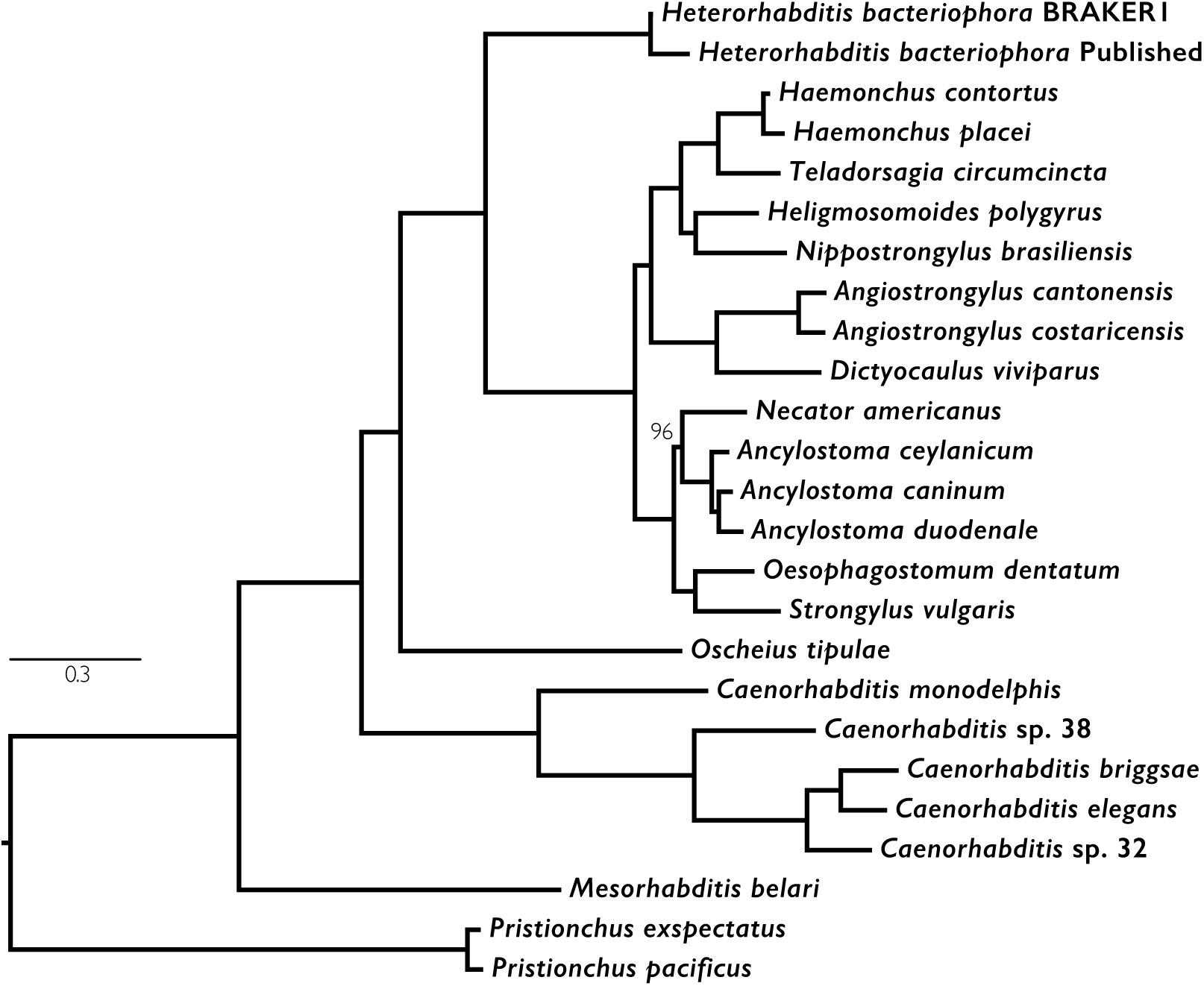
Maximum likelihood phylogeny of selected rhabditine (Clade V) nematodes. A supermatrix of aligned amino acid sequences from orthologous loci from both *H. bacteriophora* predictions and a set of 23 rhabditine (Clade V) nematodes (see Supplementary Table 3) were aligned and analysed with RaxML using a PROTGAMMAGTR amino-acid substitution model. *Pristionchus* spp. were designated as the outgroup. Bootstrap support values (100 bootstraps performed) were 100 for all branches except one.

The secretome of *H. bacteriophora* has been of particular interest as it may contain proteins involved in symbiotic interactions with *P. luminescens,* and proteins crucial to invasion and survival within the insect haemocoel. In the original publication, only 603 proteins (2.8% of the proteome) were predicted to be secreted [13]. This proportion is much lower than in free living nematodes such as *C. elegans* and it was postulated that *H. bacteriophora* relies on *P. luminescens* for secreted effectors [13]. The signal peptide detection method used in the original analyses was not described [13]. We used SignalP version 4.1 within Interproscan to annotate proteins in both the BRAKER1 and published *H. bacteriophora* proteomes. Proteins having a predicted signal peptide but no transmembrane domain were classified as secreted. We identified 1023 (6.5%) putative secreted proteins in the BRAKER1/soft-masked proteome and 1065 (5.1%) in the published proteome. By the same method other rhabditine (Clade V) nematodes that do not have known symbiotic associations with bacteria, such as *Teladorsagia circumcincta,* had comparable secretome sizes to *H. bacteriophora* (Supplementary File 8). This suggests that *H. bacteriophora* does not have a reduced secretome compared to other, related nematodes that do not have symbiont partners.

### Discussion

Assembly of, and genefinding in, new genomes is a challenging task, and especially so in larger genomes and those phylogenetically distant from any previously analysed exemplar. When applied *de novo* to datasets from extremely well-assembled and well-annotated model species, even the best methods fail to recover fully contiguous assemblies and yield predicted gene sets that have poor correspondence with the known truth [27]. A major issue with primary assemblies and gene sets arises when exceptional findings are taken at face value, and used to assert exceptional biology in a target species [28]. Where these exceptions are in fact the result of methodological failings, the scientific record, including the public databases, becomes contaminated. At best, erroneous assertions can be quickly checked and corrected, but at worst they can mislead and inhibit subsequent work.

A second concern arises from the recognition that while no method can currently produce perfect assemblies and perfect gene sets from raw data, analyses using the same toolsets will resemble each other and reflect the successes and failings of the particulars of the algorithms employed. However, when comparing genome assemblies and gene sets produced by different pipelines, it may be that the disparity in output generated by different pipelines dominates any signal from biology. Genomes assembled and annotated with the same tools will look more similar, and in a pool of assemblies and protein sets the one species that used a variant process will be flagged as exceptional. Again, the model organisms show the way: as new data and new scrutiny is added to the genome, better and better analyses are available. With additional analysis, and additional independent data, genome and gene predictions can be improved markedly for any species [29].

Here we examined the “outlier” whole-genome protein predictions from the entomopathogenic nematode *H. bacteriophora* [13]. The original publication noted that the number of novel proteins (those restricted to *H. bacteriophora)* was particularly large, while the number of secreted proteins was rather small, and suggested that these genome features might be a result of evolution to the species’ novel lifestyle (which includes an essential symbiosis with the bacterium *P. luminescens*). Overall we found that while the published genome sequence had a small amount of bacterial contamination, and a small number of “nematode” genes were predicted from these contaminants, the assembly itself was of high quality. Our re-prediction of the gene set of *H. bacteriophora* however suggested that the excess of unique genes, the lack of secreted proteins and several other surprising features of the original gene set were likely to be artefacts of the gene prediction pipeline chosen. While our gene set was by no means perfect (for example we identified an excess of single exon genes that derive from likely repetitive sequence) it had better biological completeness and credibility.

We used the RNA-seq based annotation pipeline BRAKER1 [23], not available to the authors of the original genome publication, who used JIGSAW [14] (see Supplementary File 9). While JIGSAW achieved high sensitivity and specificity at the level of nucleotide, exon and gene predictions in the nematode genome annotation assessment project, nGASP [27], direct comparison of the sensitivity and specificity of JIGSAW and BRAKER1 has not been published to the best of our knowledge. BRAKER1 has been shown to give superior prediction results over *ab initio* GeneMark-ES, or *ab initio* AUGUSTUS alone [23]. In particular, BRAKER1 is able to better use transcriptome data for gene finding. While we supplied only low volumes of Sanger-sequenced ESTs and a partial Roche 454 transcriptome to BRAKER1, the resulting gene set has much improved numerical and biological scores. In particular we note that the biological completeness of the predicted gene set now matches that of the genome sequence from which it was derived (Table 2).

The published gene set had an unusually high proportion (8.9%) of non-canonical (5’ GC – AG 3’) introns. While most genomes have a low proportion of non-canonical introns (usually approximately 0.5% of all introns), some species have markedly higher proportions [19]. The high proportion found initially in *H. bacteriophora* could perhaps have been taken as a warning that the prediction set was of concern. We note that gene predictors can be set to disallow any predictions that require non-canonical splicing, and many published genomes have zero non-canonical introns. These gene prediction sets are likely to categorically miss true non-canonically spliced genes.

The new BRAKER1 gene prediction set had many fewer species-unique genes (7.1%) than did the original (42.7%) when compared to 23 other related nematodes. We regard this reduction in novelty as indicative of a better prediction, as, for example, *C. elegans*, the best-annotated nematode genome, had only 9.2% of species unique genes in our analysis. Having a large proportion of orphan proteins is not unique to the published *H. bacteriophora* predictions. Nearly half (47%) of the gene predictions in *Pristionchus pacificus* were reported to have no homologues in fifteen other nematode species [30]. Evaluation of proteomic and transcriptomic evidence, as well as patterns of synonymous and non-synonymous substitution, suggested that as many as 42-81% of these genes were in fact expressed [31]. Therefore the high proportion of orphan genes in *H. bacteriophora* is not *prima facie* evidence of poor gene predictions. Expanded transcriptomic and comparative data are needed to build on the work we have presented in affirming the true *H. bacteriophora* gene set.

Biological pest control agents may become increasingly important for ensuring crop protection in the future [32]. A number of factors currently limit the commercial applicability of *H. bacteriophora,* including their short shelf life, susceptibility to environmental stress and limited insect tropism [12, 33]. Accurate genome annotation will assist in the analysis of *H. bacteriophora,* facilitating the exploration of genes involved in its parasitic and symbiotic interactions, and supporting genetic manipulation to enhance its utility as a biological control agent.

### Methods

#### Input data and data availability

The *H. bacteriophora* genome and annotations [13] were downloaded from Wormbase Parasite (WBPS8) [34]. ESTs [35, 36] were obtained from NCBI dbEST [37]. Roche 454 transcriptome data [13] were obtained from the Short Read Archive. *H. bacteriophora* strain Gebre, a gift from Adler Dillman, was inbred by selfing single hermaphrodites for five generations to generate the strain G2a1223. New Illumina HiSeq2000, paired end, 75 base data were generated from *H. bacteriophora* G2a1223 genomic DNA by the Millard and Muriel Jacobs Genetics and Genomics Laboratory at Caltech. They will be deposited in SRA.

The revised gene annotations for *H. bacteriophora* will be submitted to the INSDC. The Supplementary files for this manuscript are additionally available at https://github.com/DRL/mclean2017. All custom scripts developed for this manuscript are available at https://github.com/DRL/mclean2017.

#### Contaminant screening and Removal of Low Coverage Scaffolds

The assembly scaffolds were aligned to the NCBI nucleotide (nt) database, release 204, using Nucleotide-Nucleotide BLAST v2.6.0+ (RRID:SCR_008419) in megablast mode, with an e-value cut off of 1e^-25^ and a culling limit of 2 [38]. *H. bacteriophora* hits were excluded from the search using a list of all *H. bacteriophora* associated gene identifiers downloaded from NCBI GenBank nucleotide database, release 219. Raw, paired-end Illumina reads from the re-sequencing project were mapped against the assembly, as paired, using Burrows-Wheeler Aligner (BWA) v0.7.15 (RRID:SCR_010910) in mem mode with default options [39]. The output was converted to a BAM file using Samtools v1.3.1 (RRID:SCR_002105) [40] and overall mapping statistics generated in flagstat mode.

Blobtools v0.9.19 [20] was used to create taxon annotated GC-coverage plots for the published assembly, using the Nucleotide-Nucleotide BLAST and raw read mapping results. Scaffolds that did not have Nematoda as a top BLAST hit at the phylum level were identified, and the species-level top BLAST hit, length of scaffold, and scaffold mean base coverage were extracted from the Blobology output. Scaffolds with a mean base coverage of <10x were identified from the output of the Blobology pipeline and removed from the assembly. A list of excluded scaffolds is available in Supplementary File 1.

#### Generation of BRAKER1 Gene Predictions

Before annotation the published assembly was soft masked for known Nematoda repeats from the RepeatMasker Library v4.0.6 using RepeatMasker v4.0.6 (RRID:SCR_012954) [41] with default options. The two publicly available Roche 454 RNA-seq data files were adaptor and quality-trimmed using BBDuk v36.92 (unpublished toolkit from Joint Genome Institute, n.d.). Reads below an average quality of 10 or shorter than 25 nucleotides were discarded. Regions with average quality below 20 were trimmed. The cleaned reads were mapped to the soft masked assembly using STAR v2.5 (RRID:SCR_005622) with default options [42, 43]. The soft masked assembly was annotated with BRAKER1 [23] with guidance from the mapping output from STAR. An identical annotation method was applied to a hard masked version of the assembly. Hard masking was for known Nematoda repeats from the RepeatMasker Library v4.0.6 using RepeatMasker v4.0.6 with default options. The published and BRAKER1 proteomes were compared using DIAMOND v0.9.5 [25] in BLASTP mode to the Uniref90 database (release 03/2017) [24] with an expectation value cut-off of 1e^-5^ and no limit on the number of target sequences. Hits to *H. bacteriophora* proteins were removed using its TaxonID.

#### Gene Prediction Statistics

Gene-level statistical summaries were calculated including only the longest isoforms of the BRAKER1 gene predictions. The longest isoform for each gene in the BRAKER1 *H. bacteriophora* annotation was identified from the general feature format file, and then selected from the protein FASTA files. The general feature format file (GFF) for the published gene predictions did not contain any isoforms and was analysed in its entirety. Mean protein lengths were calculated from the amino-acid protein sequence files. Introns were inferred for the published GFF file using GenomeTools v1.5.9 in -addintrons mode [44]. Intron frequencies were then calculated for the published and BRAKER1 annotations from their respective GFF files. Exon frequencies were calculated for the published annotations directly from the GFF file. For the BRAKER1 annotations, exon frequency per gene was assumed to be equivalent to coding DNA sequence (CDS) frequency and inferred from the general feature format file as exon features were not included in the GFF. Intron frequency histograms and bar plots were generated in Rstudio v1.0.136 (RRID:SCR_005622) with R v3.3.2 (RRID:SCR_001905) and in some instances the package ggplot2 v2.2.1. As intron frequency lists did not contain single exon genes (those with no introns), these were added manually to the intron frequency lists in Microsoft Excel before importing the data into Rstudio.

The proportion of introns with GC – AG splice junctions was assessed for the gene models of *C. elegans* (WS258), and the published and BRAKER1/soft-masked gene models of *H. bacteriophora.* Intronic features were added to GFF3 files using GenomeTools v1.5.9 [44] (‘gt gff3 -sort -tidy -retainids -fixregionboundaries -addintrons’) and and splice sites were extracted using the script extractRegionFromCoordinates.py [19]. Results were visualised using the script plot_GCAG_counts.R (see https://github.com/DRL/mclean2017).

Gene features, extracted from the GFF files, were assessed for overlap using bedtools v2.26 (RRID:SCR_006646) in intersect mode [45]. Only genes on the same strand were considered to be overlapping. To calculate the number of identical proteins shared between the published and BRAKER1 proteomes non-redundant protein fasta files were generated using cd-hit v4.6.1 (RRID:SCR_007105) [46] for the BRAKER1 and published predictions. The files were concatenated, sorted and unique sequences counted using unix command line tools.

BUSCO v2.0.1 (RRID:SCR_015008) [18], with Eukaryota as the lineage dataset, and *Caenorhabditis* as the species parameter for orthologue finding was applied to both proteomes and the published assembly to calculate BUSCO scores. CEGMA (RRID:SCR_015055) [17] was run on the published genome sequence. BWA was used with default settings to map the RNA-seq datasets to the CDS transcripts from the published and BRAKER1 annotations and the summary statistics obtained with Samtools v1.3.1 in flagstat mode.

#### Protein orthology analyses

OrthoFinder v1.1.4 [26] with default settings was used to identify orthologous groups in the proteomes of 23 Clade V nematodes with the addition of either the BRAKER1/soft-masked and published *H. bacteriophora* proteomes separately or simultaneously. The proteomes for the 23 Clade V nematodes were downloaded from WBPS8 (available at: http://parasite.wormbase.org/index.html) or GenomeHubs.org (available at http://ensembl.caenorhabditis.org/index.html), and detailed source information is available in Supplementary File 5. All proteomes were filtered to contain only the longest isoform of each gene, and for all proteomes (except the BRAKER1/soft-masked *H. bacteriophora* protein set), proteins less than 30 amino-acids in length were excluded before clustering. For the *H. bacteriophora* BRAKER1/soft-masked protein set, proteins less than 30 amino-acids (SF5.2) were removed manually from the orthofinder clustering statistics after clustering. None of these proteins seeded new clusters and are therefore will not have influenced the clustering results. Kinfin v0.9 [47], was used with default settings to identify true and fuzzy 1-to-1 orthologues, and their associated species specific statistics. Fuzzy 1-to-1 orthologues are true 1-to-1 orthologues for greater than 75% of the species clustered. For the clustering analysis presented in Supplementary File 3, the BRAKER1/soft masked and published proteomes were clustered simultaneously to the 23 other Clade V nematode proteomes, and singletons, and species-specific clusters were excluded.

#### Interproscan and search for transposons

Interproscan v5.19-58.0 (RRID:SCR_005829) [48] was used in protein mode to identify matches with the BRAKER1 and published *H. bacteriophora* predicted proteomes in the following databases: TIGRFAM v15.0, ProDom v2006.1, SMART-7.1, SignalP-EUK v4.1, PrositePatterns v20.119. PRINTS v42.0, SuperFamily v1.75, Pfam v29.0, and PrositeProfiles v20. 119. InterProScan was run with the option for all match calculations to be run locally and with gene ontology annotation activated. The number of single exon genes with similarity to transposons or transposases in the BRAKER1/soft masked predictions was calculated by searching the full InterProScan results for the strings ‘Transposon’, ‘transposon’, ‘Transposase’, or ‘transposase’ and the number of single exon gene InterProScan results containing these terms counted. InterProScan results from searching the SignalP-EUK-4.1 database were queried to identify putative secreted proteins. Those with a predicted signal peptide but no transmembrane region were considered to be secreted.

#### Phylogenetic Analyses

Both *H. bacteriophora* proteomes were clustered simultaneously with the 23 Clade V nematode proteomes into orthologous groups using Orthofinder v1.0 [26]. The fuzzy 1-to-1 orthologues were extracted and processed using GNU parallel [49]. They were aligned using MAFFT v7.267 (RR1D:SCR 011811) [50], and the alignments trimmed with NOISY v1.5.12. A maximum likelihood gene tree was generated for each orthologue using RaXML v8.1.20 (RRID:SCR_006086) [51] with a PROTGAMMAGTR amino-acid substitution model. Rapid Bootstrap analysis and search for the best -scoring ML tree within one program run with 100 rapid bootstrap replicates was used. The trees were pruned using PhyloTreePruner v1.0 [52] to remove paralogues, with 0.5 as the bootstrap cutoff and a minimum of 20 species in the orthogroup after pruning for inclusion in the supermatrix. Where species had more than one putative orthologue in an orthogroup the longest was selected. The remaining 897 orthogroups were re-aligned using MAFFT v7.267, trimmed with NOISY v1.5.12 and concatenated into a supermatrix using FASconCAT v.1.0 [53]. A supermatrix maximum-likelihood tree was generated using RAxML with the rapid hill climbing algorithm (default), with a PROTGAMMAGTR amino-acid substitution model and 100 bootstrap replicates. *Pristionchus* spp. were designated as the outgroup. The tree was visualised in Dendroscope v3.5.9 [54].

## Acknowledgements

This project was supported by FMs Wellcome Trust-funded graduate programme [204052/Z/16/Z]. Sujai Kumar, Lewis Stevens, Carlos Caurcel and Elisabeth Sjokvist offered expert technical support and advice. Igor Antoshechkin of the Millard and Muriel Jacobs Genetics and Genomics Laboratory at Caltech assisted with Illumina sequencing. Adler Dillman provided the parental strain for the inbred *H. bacteriophora* strain G2a1223.

## Supplementary Files

The supplementary files for this work are described below. All Supplementary files are available at https://github.com/DRL/mclean2017.

### Supplementary file 1: Scaffolds and contigs removed from the *Heterorhabditis bacteriophora* assembly because of low coverage in the new whole genome sequencing dataset

Text file.

### Supplementary File 2: BRAKER1/soft-masked annotations of *Heterorhabditis bacteriophora*

A zipped archive (14.1 Mb) of the BRAKER1/soft-masked annotations of *Heterorhabditis bacteriophora*. The archive contains three text files: the GFF format file, the GTF format file and the amino acid sequences of the protein predictions in FASTA format.

### Supplementary File 3: BRAKER1/hard-masked annotations of *Heterorhabditis bacteriophora.*

A zipped archive (13.4 Mb) of the BRAKER1/hard-masked annotations of *Heterorhabditis bacteriophora.* The archive contains three text files: the GFF format file, the GTF format file and the amino acid sequences of the protein predictions in FASTA format.

### Supplementary File 4: Comparison of the BRAKER1/soft-masked and BRAKER1/hard-masked gene predictions from *Heterorhabditis bacteriophora*

Tab-delimited text file.

### Supplementary File 5: OrthoFinder analyses of *Heterorhabditis bacteriophora* predicted proteomes

A zipped archive (20.3 Mb) of the OrthoFinder analyses of *Heterorhabditis bacteriophora* predicted proteomes with 23 other nematode species. The archive contains the following files:

SF5.1 A list of the proteomes included in the OrthoFinder analyses (text format file)

SF5.2 List of *Heterorhabditis bacteriophora* proteins of length <30 amino acids excluded from the OrthoFinder analyses (text format file).

SF5.3 The OrthoFinder output files. A zipped archive of the three OrthoFinder clustering result files (published *H. bacteriophora* + 23 species; BRAKER1/soft-masked + 23 species: published + soft-masked + 23 species).

SF5.4 Table with count of orthogroups at each contribution ratio from the BRAKER1/soft-masked and published proteomes after clustering with 23 other Clade V nematodes.

### Supplementary File 6: KinFin analyses of *Heterorhabditis bacteriophora* predicted proteomes

A zipped archive (27.8 Mb) of the KinFin analyses from the OrthoFinder analyses of *Heterorhabditis bacteriophora* predicted proteomes.

### Supplementary File 7: Phylogenetic analyses of *Heterorhabditis bacteriophora* predicted proteomes

A zipped archive (11.2 Mb) of the supermatrix alignment and the phylogenetic trees produced for the the analyses of the *Heterorhabditis bacteriophora* proteomes. The archive contains the following files:

SF7.1 Alignments of orthogroups used to build the supermatrix (directory of aligned sequences in fasta format).

SF7.2 Supermatrix of aligned sequences (FASTA .fas format file).

SF7.3 Phylogenetic analysis output files (NEWICK format text file).

### Supplementary File 8: Secretome analysis of *Heterorhabditis bacteriophora* predicted proteomes

Secretome analyses of *Heterorhabditis bacteriophora* and other rhabditine nematodes. The zipped archive (8 kb) contains the following text format files.

SF8.1 Secretome predictions from the published Bai *et al.* (2013) protein predictions.

SF8.2 Secretome predictions from the BRAKER1/soft-masked predictions.

### Supplementary File 9: BRAKER1 and JIGSAW annotation pipelines

Figure illustrating the differences between the BRAKER1 and the Bai et al 2013 JIGSAW prediction methods used for *Heterorhabditis bacteriophora*. PDF file.

